# Learning Divisive Normalization in Primary Visual Cortex

**DOI:** 10.1101/767285

**Authors:** Max F. Burg, Santiago A. Cadena, George H. Denfield, Edgar Y. Walker, Andreas S. Tolias, Matthias Bethge, Alexander S. Ecker

## Abstract

Divisive normalization (DN) is a prominent computational building block in the brain that has been proposed as a canonical cortical operation. Numerous experimental studies have verified its importance for capturing nonlinear neural response properties to simple, artificial stimuli, and computational studies suggest that DN is also an important component for processing natural stimuli. However, we lack quantitative models of DN that are directly informed by measurements of spiking responses in the brain and applicable to arbitrary stimuli. Here, we propose a DN model that is applicable to arbitrary input images. We test its ability to predict how neurons in macaque primary visual cortex (V1) respond to natural images, with a focus on nonlinear response properties within the classical receptive field. Our model consists of one layer of subunits followed by learned orientation-specific DN. It outperforms linear-nonlinear and wavelet-based feature representations and makes a significant step towards the performance of state-of-the-art convolutional neural network (CNN) models. Unlike deep CNNs, our compact DN model offers a direct interpretation of the nature of normalization. By inspecting the learned normalization pool of our model, we gained insights into a long-standing question about the tuning properties of DN that update the current textbook description: we found that within the receptive field oriented features were normalized preferentially by features with similar orientation rather than non-specifically as currently assumed.

**Author summary:** Divisive normalization (DN) is a computational building block throughout sensory processing in the brain. We currently lack an understanding of what role this normalization mechanism plays when processing complex stimuli like natural images. Here, we use modern machine learning methods to build a general DN model that is directly informed by data from primary visual cortex (V1). Contrary to high-predictive deep learning models, our DN-based model’s parameters offer a straightforward interpretation of the nature of normalization. Within the receptive field, we found that neurons responding strongly to a specific orientation are preferentially normalized by other neurons that are highly active for similar orientations, rather than being normalized by all neurons as it is currently assumed by textbook models.

## 1 Introduction

A crucial step towards understanding the visual system is to build models that predict neural responses to arbitrary stimuli with high accuracy (Carandini et al., 2005). The classical standard models of primary visual cortex (V1) are based on linear-nonlinear models (Simoncelli et al., 2004), energy models (Adelson and Bergen, 1985) and subunit (LN-LN) models (Rust et al., 2005; Touryan et al., 2005; Willmore et al., 2008; Butts et al., 2011; McFarland et al., 2013; Vintch et al., 2015). Fueled by advances in machine learning technology, recent studies have shown that multi-layer convolutional neural networks (CNNs) can significantly improve the prediction of neural responses to complex images and videos at several stages of the visual pathway, outperforming classical models (Yamins et al., 2014; Khaligh-Razavi and Kriegeskorte, 2014; McIntosh et al., 2016; Zhang et al., 2019; Cadena et al., 2019; Kindel et al., 2019; Walker et al., 2019; Sinz et al., 2018). The current state-of-the-art data-driven model of single-unit activity in monkey V1 is a three-layer CNN (Cadena et al., 2019). However, it is challenging to gain insights into V1 function from the features produced by deeper layers of such models. In particular, we do not have first principles explaining the kind of nonlinearities approximated in successive layers of CNNs, or if these nonlinearities can be described in a compact way in the first place.

A promising candidate to facilitate a more succinct description of V1 neurons is to replace the computations from the CNN’s deeper layers by a divisive normalization nonlinearity (Heeger, 1992) that is easier to understand. Divisive normalization has been proposed as a canonical neural computation present throughout the visual pathway (Carandini and Heeger, 2012) because it (1) explains a wide variety of neurophysiological phenomena using simple stimuli (Carandini and Heeger, 2012; Sawada and Petrov, 2017) and (2) can be derived from first principles of redundancy reduction (Schwartz and Simoncelli, 2001; Sinz and Bethge, 2008). The significance of DN is also supported by its recent success in computer vision where it enables state-of-the-art natural image compression with high perceptual quality (Ballé et al., 2017).

Divisive normalization has been invoked to explain several nonlinear neurophysiological phenomena in V1, which can be classified into (1) phenomena that are restricted to the receptive field (RF) only or (2) phenomena that involve an interaction between receptive field and its surround. A prominent example for a mechanism restricted to the receptive field is cross-orientation inhibition (Bonds, 1989; Morrone et al., 1982; DeAngelis et al., 1992; Heeger, 1992; Carandini et al., 1997; Freeman et al., 2002; Busse et al., 2009), whereas an example for an interaction of the RF with its spatially adjacent surround is surround suppression (Blakemore and Tobin, 1972; DeAngelis et al., 1994; Cavanaugh et al., 2002a,b; Coen-Cagli et al., 2015). In our work, we focus on the first aspect: divisive normalization *within* the RF and cross-orientation inhibition. In this phenomenon, the response of a neuron to a driving grating stimulus in the RF is suppressed by superimposing a second grating also within the RF that does not elicit a response when presented alone: for instance, a grating with orientation orthogonal to the neuron’s preferred orientation.

The basic idea of divisive normalization (Fig. 1A) is that a neuron’s driving input is normalized divisively by a weighted sum over nearby neurons’ responses (Heeger, 1992; Carandini and Heeger, 2012). While the general idea is simple, elegant and powerful, our current knowledge of DN is limited in two important ways: (1) DN has been studied mostly with simple, synthetic stimuli, but its implications under natural image stimulation are unclear. For example we do not know whether incorporating DN into system identification models predicting neural responses to natural stimuli improves performance. (2) We currently do not know how response properties of neurons with overlapping receptive fields determine whether a neuron contributes to the normalization pool of another neuron and, if so, with which strength (Carandini et al., 1997).

**Figure 1:**
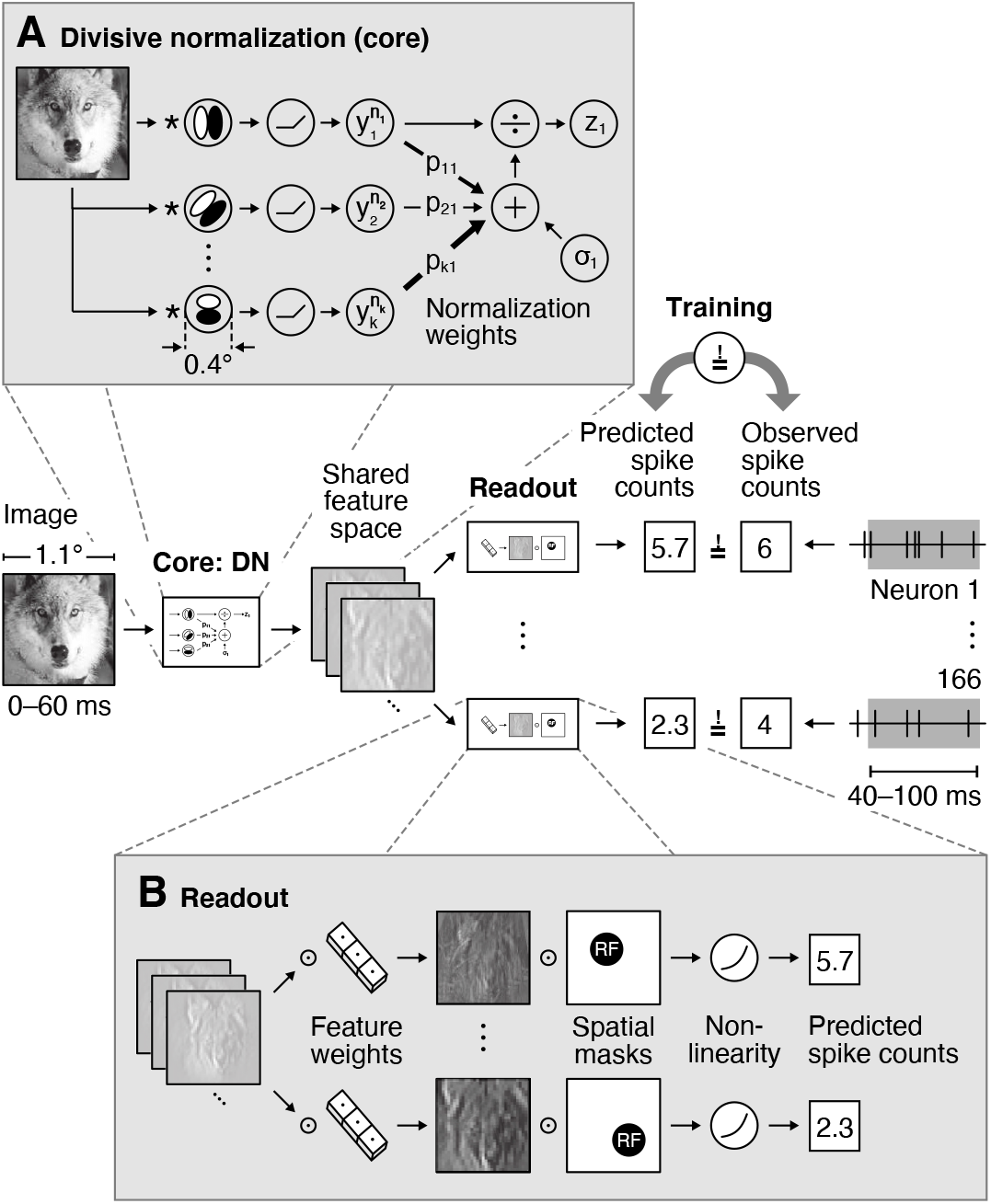
Overview of our divisive normalization (DN) model. The model takes as input an image covering 1.1°of visual field and predicts neurons’ spike counts in response to this image (details in Fig. 2). The model is split into two parts: a *core* that computes a shared nonlinear feature space and a *readout* that maps the shared feature space individually to each neuron’s spike count. **A**. Divisive normalization mechanism (simplified). The visual input is convolved with 32 filters covering 0.4°of visual field and then rectified and exponentiated to produce an excitatory output. The output of each filter is then divided by a weighted sum of the excitatory outputs of all filters with normalization weights *p*_*kl*_ and a semi-saturation constant *σ*_*l*_. In our general formulation, all weights and constants are learned from the data. **B**. Readout that maps the shared feature space to each neuron’s spike count through an individual weighted sum over the entire shared feature space and a pointwise output non-linearity. The readout weights are factorized into a feature vector – capturing the nonlinear feature(s) that a neuron computes – and a spatial mask – localizing each neuron’s receptive field (RF).

To explain normalization phenomena within the receptive field like cross-orientation inhibition, current models of divisive normalization assume that all nearby neurons with diverse orientation tuning preferences and with similar receptive field locations contribute equally to the normalization pool (Heeger, 1992; Carandini et al., 1997; Busse et al., 2009). However, some earlier experimental studies suggest that this assumption may not be correct for some neurons (Bonds, 1989; DeAngelis et al., 1992), and normative models of normalization predict that the magnitude with which a given neuron contributes to another neuron’s normalization should depend on the relationship of their response properties (Schwartz and Simoncelli, 2001).

In this paper, we address the two main questions raised above: (1) How well does a data-driven model with divisive normalization predict V1 responses to natural images, compared to classical subunit models and deep convolutional neural networks, and (2) what stimulus features are learned for normalizing a V1 neuron’s response in relation to its favoured feature selectivity within the receptive field?

We focused on effects restricted to the receptive field and on models that only account for local normalization interactions between neurons with overlapping receptive fields. We developed an end-to-end trainable model predicting V1 spike counts from natural stimuli, replacing the CNNs deeper layers by the canonical neural DN computation (Carandini and Heeger, 2012) and learning the filters of all neurons as well as their normalization weights directly from the data. We also explored how our DN model could be extended in a way that might enable it to capture surround interactions from outside of the RF, however, our control experiments demonstrate that our results are very unlikely to include the extra-classical surround.

We fitted our DN model to monkey V1 responses to natural images and found that it outperforms linear-nonlinear and subunit models, suggesting that DN mechanisms are useful for capturing nonlinear computations unlocked by natural stimuli beyond those evoked by simpler gratings. The power of our model was further strengthened as it captured cross-orientation inhibition within the receptive field, brought to light via *in silico* experiments. In contrast to the three-layer CNN, which also reproduced this phenomenon, the single divisive layer of the DN model provides direct interpretations of the normalization pool. These revealed that oriented features were preferentially normalized by other features with similar orientation, which quantitatively improved predictive performance over a model with nonspecific normalization. However, the three-layer CNN outperformed our DN models, suggesting that there are additional non-linear mechanisms beyond the ones captured by our DN model (e. g. phase invariance and divisive effects), that are required to achieve optimal performance in modeling V1 responses. Thus, our work advances our understanding of V1 function, highlighting the importance of orientation-specific gain control to predict responses to natural images, and shows how embedding theories of neural processing into the architectures of generic data-driven models can improve our understanding of sensory processing in the brain.

## 2 Results

### 2.1 Learning divisive normalization

The basic idea of divisive normalization (Fig. 1A) is that the response of neuron *l*

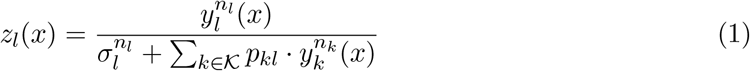

is given by its driving input activity *y*_*l*_(*x*) exponentiated by *n*_*l*_ and divisively normalized by a weighted sum over nearby neurons’ exponentiated responses 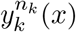 (*x*) (Heeger, 1992; Carandini and Heeger, 2012), where *x* represents the stimulus, *σ*_*l*_ is a semi-saturation constant, and all quantities are non-negative. Here, the set of normalizing neurons 𝒦 and the normalization weights *p*_*kl*_ define which neurons *k* contribute to the normalization pool of neuron *l* and with what strength, respectively.

While this formulation is straightforward to write down, it is challenging to build quantitative models based on it that are applicable to arbitrary inputs. The denominator depends on a potentially large population of neurons – which is unknown in general – and the structure of the normalization weights has been studied only using very restricted sets of simple stimuli such as oriented gratings and bars. Previous modeling work on divisive normalization has therefore made specific assumptions about the filter properties of the underlying neuronal population and either modeled only a closed set of stimuli such as gratings of different orientation (Heeger, 1992; Carandini et al., 1997; Freeman et al., 2002; Heuer and Britten, 2002; Busse et al., 2009) or evaluated models only qualitatively (Schwartz and Simoncelli, 2001; Wainwright et al., 2002; Froudarakis et al., 2014).

We developed a general, image-computable predictive model of divisive normalization following Eq. (1), which is applicable to arbitrary images and whose parameters are learned by optimizing the accuracy of the model in predicting the spiking activity of a large number of neurons in response to natural images (see Fig. 1). Our model builds on a recent innovation in predictive modeling (Antolík et al., 2016; Klindt et al., 2017; Batty et al., 2017; McIntosh et al., 2016; Cadena et al., 2019), jointly modeling all recorded neurons instead of learning a predictive model for each neuron individually. Because many neurons perform similar computations – up to shifts in receptive field location – jointly modeling them makes more efficient use of the data and we can learn more complex models. The basic idea is to split the model into two parts (Fig. 1): (A) a *core* that transforms the input image into nonlinear features shared among all neurons, and (B) a *readout* that linearly combines the features to produce a prediction of each neuron’s response.

We use a convolutional network for the core, whose architecture lends itself very well to model divisive normalization. By construction, we have a model that contains all filters necessary to account for the recorded neurons’ responses. All of these filter responses are automatically extracted at each location, providing a good approximation of the underlying population of neurons in the brain although it is only sparsely sampled during the experiment. As a consequence, we can optimize the pool of neurons providing normalizing inputs and their corresponding weights *p*_*kl*_ (Eq. 1) to account for the neural responses. In this work we mostly focus on normalization effects from neurons with overlapping receptive fields, rather than investigating interactions with the RFs’ surround.

In summary, our model’s core (Fig. 1A) consists of a set of convolutional filters *w*_*l*_ and biases *o*_*l*_ (we use 32) that provide the driving inputs *y*_*l*_ from an image *x* at all locations in the neurons’ RFs,

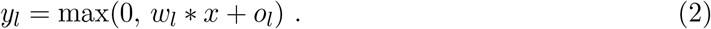

As a specific feature is extracted by its according convolutional filter at each spatial location of the image, the driving responses *y*_*l*_ form a map of responses for the according features, hence named feature map. This operation is followed by a DN stage (Eq. 1) in which all operations are performed element-wise across space. This core is shared among all neurons and converts the image into a set of feature maps *z*_*l*_ containing information from the neurons’ RFs. We mapped those normalized feature maps into response predictions 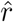 by a linear readout step (Fig. 1B) that picks the relevant features by readout weights *b* and spatial locations by readout weights *a* for each neuron *i*, followed by a pointwise output nonlinearity *ϕ*,

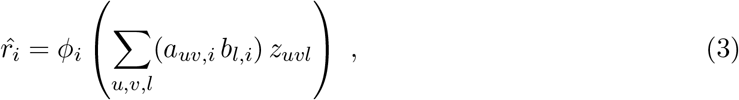

where *u, v* index space (see Methods for additional details). To ensure that the readout does not model any inhibitory or complex interactions, we constrained its weights *a, b* to be non-negative and added a sparsity prior. The non-negativity ensures that activations can only add, preventing the readout stage from accounting for any suppressive effects. The readout can, however, account for response invariances such as phase invariance of complex cells; see Methods for an in-depth explanation. While our model reflects the general formulation of divisive normalization, mathematically it is not identical to the classical formulation of DN due to the additional linear-nonlinear readout after the model’s core.

### 2.2 DN model outperforms subunit model, half-way closing the gap to state-of-the-art performance

We fitted the model described above to a dataset of 166 neurons recorded in V1 of two awake, fixating monkeys (data from Cadena et al. 2019), who viewed a fast sequence of localized natural images and textures (Fig. 2). The stimuli were placed at the neurons’ receptive fields and primarily stimulated the center (i. e. the RF) as they covered one degree of visual field, quickly fading off reaching zero contrast at two degrees (see Discussion and Methods for additional details). Images were shown for 60 ms each, without blanks in between. Single unit activity was recorded with laminar silicon probes sampling from all cortical layers. We fit the model jointly to the responses of all neurons. As neurons were recorded in 17 recording sessions, the dataset sampled a diverse range of preferred orientations. The objective function during training was to minimize the difference between the model’s prediction and the observed spike counts of the neurons in a time window 40–100 ms after image onset (to account for response latency).

**Figure 2:**
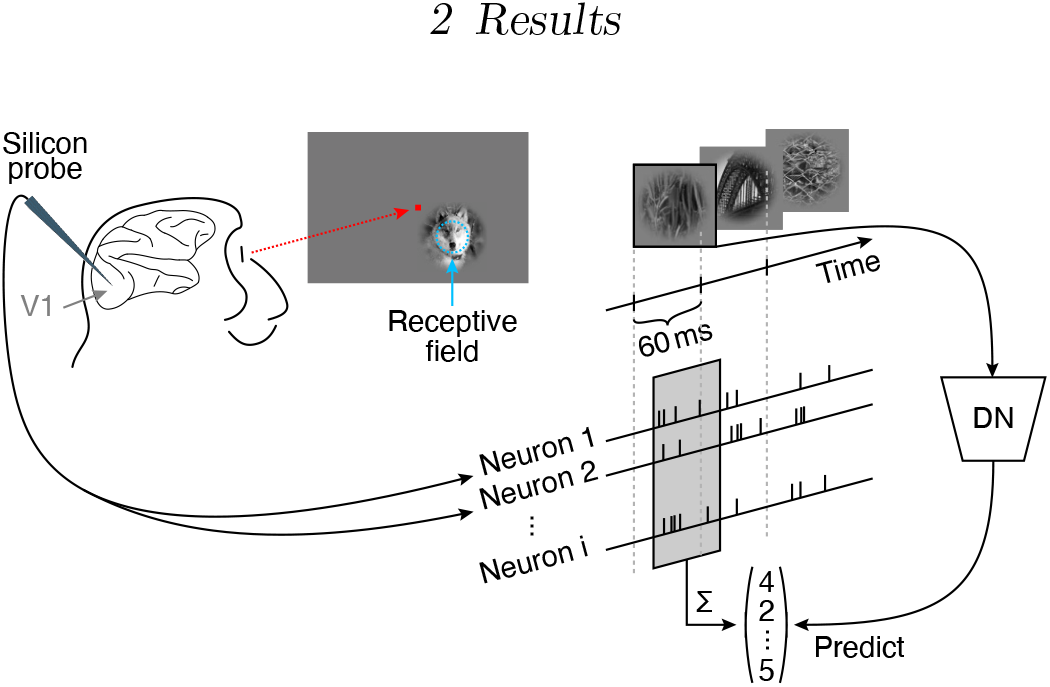
Experimental paradigm from Cadena et al. (2019). Natural images were flashed to a monkey covering 2°of their visual angle, and located at the center of the multi-unit receptive field. Multiple neurons were isolated from recordings with silicon probes inserted into V1 (Denfield et al., 2018). Natural images were shown in a fast sequence without blanks, each presented for 60 ms. Spike counts from all isolated neurons corresponding to each image were extracted from a window 40 ms after the image onset lasting 60 ms.

To evaluate model performance, we estimated the fraction of explainable variance explained (FEV), which quantifies the fraction of the stimulus-driven response variance that is accounted for by the model, and ignores unexplainable trial-to-trial variability in the response of the neurons (see Methods). A perfect model would reach a FEV of 100%. Since model performance depends on the parameters’ initial values before optimization, we initialized all weights randomly before model training and repeated this procedure multiple times. To ensure that the reported performance is not a spurious characteristic of one particular model, we picked the best ten models of a model class (assessed in terms of validation set accuracy) and report the mean performance together with its 95% confidence interval. We perform this analysis for each of the four model classes.

Subunit models are an established approach to model primary visual cortex responses (Rust et al., 2005; Touryan et al., 2005; Willmore et al., 2008; Butts et al., 2011; McFarland et al., 2013; Vintch et al., 2015). In addition to capturing a fair portion of the explainable variance (Cadena et al., 2019), their computations are easily understandable because they are rather shallow models consisting of a first stage of rectified linear filtering followed by a static nonlinearity, and then a pooling stage with a subsequent output nonlinearity. Although the final output nonlinearity is capable to learn various pointwise functions like sigmoids, in most cases (across all model types) it leads to only slight expansive deviations of the identity function. Structurally, the subunit model is the same as our DN model, but without the normalization stage. Therefore, we considered a convolutional subunit model as a strong baseline for our model. This subunit model accounted for (46.6 ± 0.2) % FEV (mean and 95% confidence interval over the ten best models selected in terms of validation set accuracy). In comparison, a regularized linear nonlinear Poisson model (LNP) only accounted for 16.3% FEV on the same dataset (Cadena et al., 2019) due to its inability to model complex cells.

As recent developments in machine learning technology have allowed us to improve predictive performance, we used the current best data-driven model as a gold standard. This model is a convolutional neural network with three convolutional layers and a linear-nonlinear readout (Cadena et al., 2019). To enable fair comparison of this model to all other models in this study, we optimized the CNN comparable to our DN model, reaching (52.3 ± 0.1) % FEV (mean and 95% confidence interval over the ten best models selected in terms of validation set accuracy; outperforming previously reported performance of 49.8% FEV; Cadena et al. 2019). However, although this CNN model outperforms the simpler subunit model, it is of higher complexity requiring a higher number of parameters.

To evaluate how well our DN model accounts for the data, we placed its performance on a scale between 0% (baseline: subunit model) and 100% (gold standard: CNN). On that scale, our DN model achieved a score of (52±3) % between the baseline and gold standard (Fig. 3; (49.6±0.1) % FEV; mean and 95% confidence interval over the ten best models selected in terms of validation set accuracy). The performance differences between all model pairs were statistically significant (pairwise Wilcoxon signed rank test on best models in terms of validation accuracy: *p* < 0.024, *N* = 166 neurons, family-wise error rate *α* = 0.05 using Holm-Bonferroni correction to account for multiple comparisons). Notably, we achieved this performance gain by simply adding the trainable DN stage to the convolutional subunit model, which quantifies the importance of divisive normalization as computational mechanism to predict neural responses in V1 under stimulation with complex, natural images. At the same time, the CNN outperformed the DN model, suggesting that further non-linear dependencies in addition to phase invariance (complex cells) and divisive normalization are required to reach optimal predictive performance.

**Figure 3:**
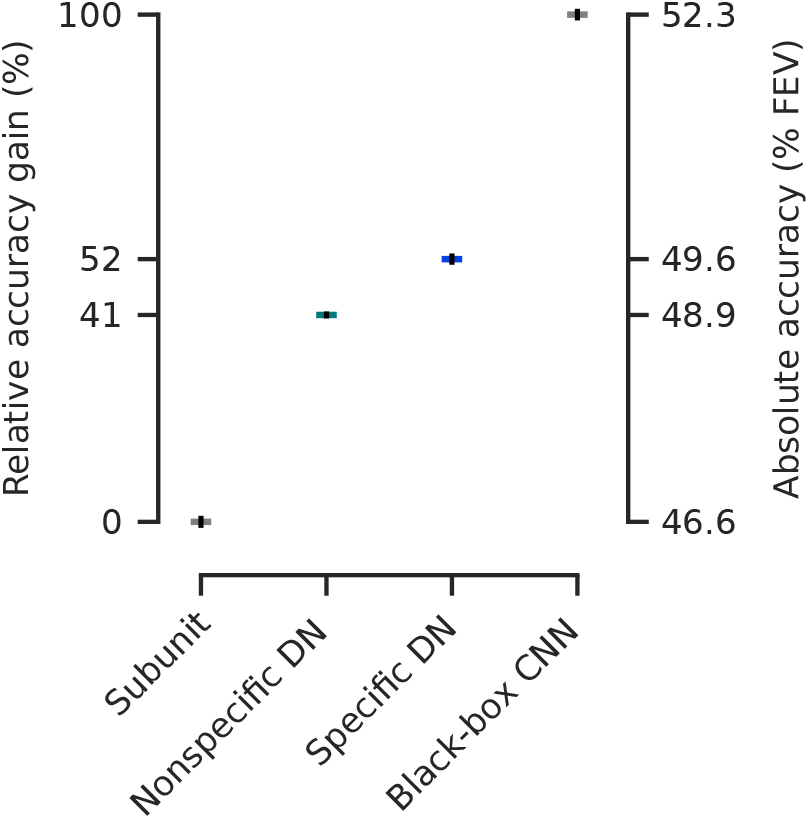
Performance comparison of our models fitted to the data from Cadena et al. (2019) relative to the gap between the best shallow model – a subunit one layer convolutional neural network (CNN) – and the deeper data-driven state-of-the-art three-layer CNN (Cadena et al., 2019). Non-specific divisive normalization (DN) accounts for 41 % of this gap, while specific DN improves it up to 52 %. Absolute values in terms of percentage of explainable variance explained (FEV) on the right (mean over the ten best models selected in terms of validation set accuracy, see main text for details). Error-bars (black) indicate the standard error of the mean. Model performance is significantly different between each model class (pairwise Wilcoxon signed rank test on best models in terms of validation accuracy: *p* < 0.024, *N* = 166 neurons, family-wise error rate *α* = 0.05 using Holm-Bonferroni correction).

### 2.3 Divisive normalization and CNN models learn cross-orientation inhibition

We wanted to test if our DN model captured non-linear interactions between the neurons inside the receptive field leading to cross-orientation inhibition, a phenomenon that was explained by DN within the RF before (Bonds, 1989; Morrone et al., 1982; DeAngelis et al., 1992; Heeger, 1992; Carandini et al., 1997; Freeman et al., 2002; Busse et al., 2009). As the gold standard three-layer convolutional neural network explains more variance than the divisive normalization model, we asked if the CNN captured cross-orientation inhibition, too. To assess if our models learned cross-orientation inhibition, we performed the experiments done before by experimentalists *in silico*. Specifically, for each unit we determined the optimal Gabor maximizing our models’ predictions. Subsequently, we combined the optimal Gabor by adding the same Gabor pattern with orthogonal orientation, obtaining a plaid stimulus (Fig. 4A insets). We then presented the plaids to our models varying the contrasts of each Gabor component and measured the resulting tuning curve for all models *in silico*, averaging across the orthogonal Gabor’s phase to get independent of any phase dependency. For the DN model and the CNN, we found cross-orientation inhibition in a number of cells: the response elicited by presenting the optimal Gabor was inhibited by increasing the contrast of the orthogonal Gabor component (Fig. 4A). However, such behavior was not clearly visible for the subunit model.

**Figure 4:**
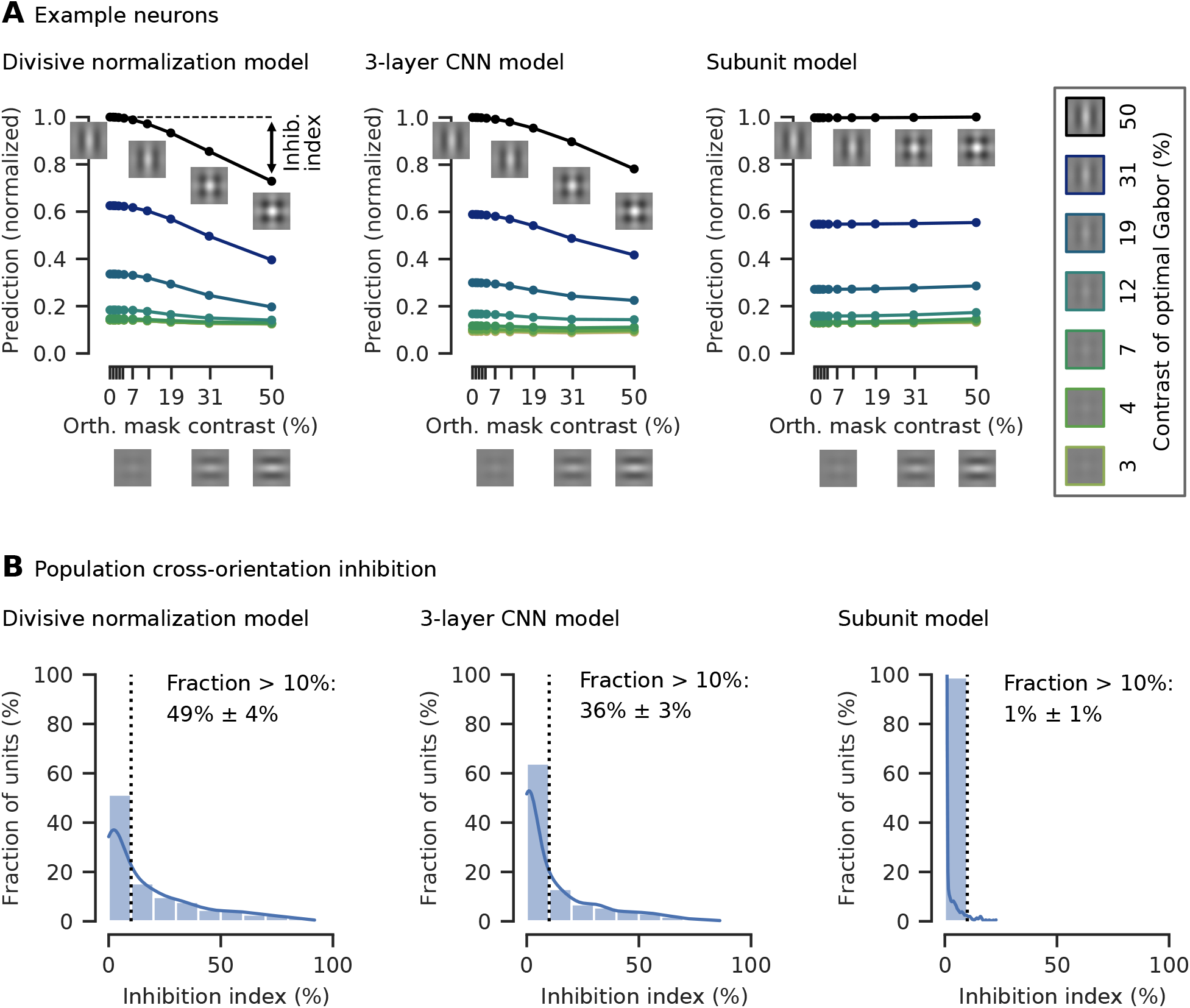
Cross-orientation inhibition was learned by our divisive normalization model and the three-layer CNN, but not significantly by the subunit model. **A**. Tuning curves for an example neuron of all three models and various contrast combinations of the optimal Gabor (box on the right, examples for contrasts of 0%, 1% and 2% not shown) and an orthogonal Gabor masking. As the contrast of the orthogonal mask increases, the model prediction (normalized by the maximum response) decreases. The cross-orientation inhibition (inhib.) index measures the percentage of response inhibition by adding the masking compared to the optimal Gabor presented alone, in this case approximately 20%. Insets: illustration of plaid stimuli, created by overlaying an optimally oriented Gabor with an orthogonal mask. **B**. Histograms of the cross-orientation inhibition indices accumulated across the best ten models (in terms of validation set accuracy) per model type, with kernel density estimate of the underlying distribution. The fraction of cells that show more than 10% cross-orientation inhibition is displayed right of the dotted line (mean and 95% confidence interval over the ten best models selected in terms of validation set accuracy). For the DN model, more cells show cross-orientation inhibition compared to the other models. The subunit model shows almost no cross-orientation inhibition.

To quantify the models’ capability to learn cross-orientation inhibition, we defined a cross-orientation inhibition index measuring the percentage an orthogonal Gabor inhibits response when added to an optimal Gabor presented alone (Fig. 4A left; see Methods for details). To determine how many cells are cross-orientation inhibited by at least 10%, we assess the highest cross-orientation inhibition index across all contrast combinations of the two component Gabors. To ensure we report a general property of all models, we quantified cross-orientation inhibition across the ten best models of each model-type (assessed in terms of performance on the validation set, only differing in the initialization of the model parameters before training). In the subunit model we found no significant cross-orientation inhibition (Fig. 4B). Although the 3% FEV performance gain of the DN model over the subunit model might seem small, the phenomenological difference between those models was quite substantial, as in the DN model half of the cells showed cross-orientation inhibition. The gold-standard three-layer CNN captured cross-orientation inhibition in fewer cells, likely approximating divisive normalization through its deeper layers as its filters in the first convolutional layer are qualitatively similar to the DN model, showing Gabor-like and blob patterns. At the same time, the CNN seems to capture additional effects from the data that are not explained by DN in the RF center, leading to the CNN’s higher performance.

### 2.4 Normalization within the receptive field is feature-specific

Having established that both the DN and the CNN models learned a phenomenon explained by divisive normalization within the RF, namely cross-orientation inhibition, we next asked which features and how strongly they contribute to the normalization pool. In the CNN model there is no explicit notion of a normalization pool that we could comprehensively analyze. On the other hand, our DN model offers a way to investigate the structure of the normalizing input within the receptive field, i. e. the image-averaged products 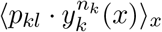 in the denominator of Eq. (1), to quantify the strength by which different features contribute to it (see Methods for details). When visualizing the feed-forward features learned by the DN model we found that several exhibited clear orientation selectivity (i. e. features with an anisotropic structure; see Fig. 5A). Thus, for these features we first wondered about the relationship between their preferred orientation and the orientation of the features by which they are preferentially normalized. To this end, we visualized the feature pair-wise normalizing input (Fig. 5A) and found that oriented features are normalized preferentially by features with similar orientation preference, while orthogonal features seem to contribute less.

**Figure 5:**
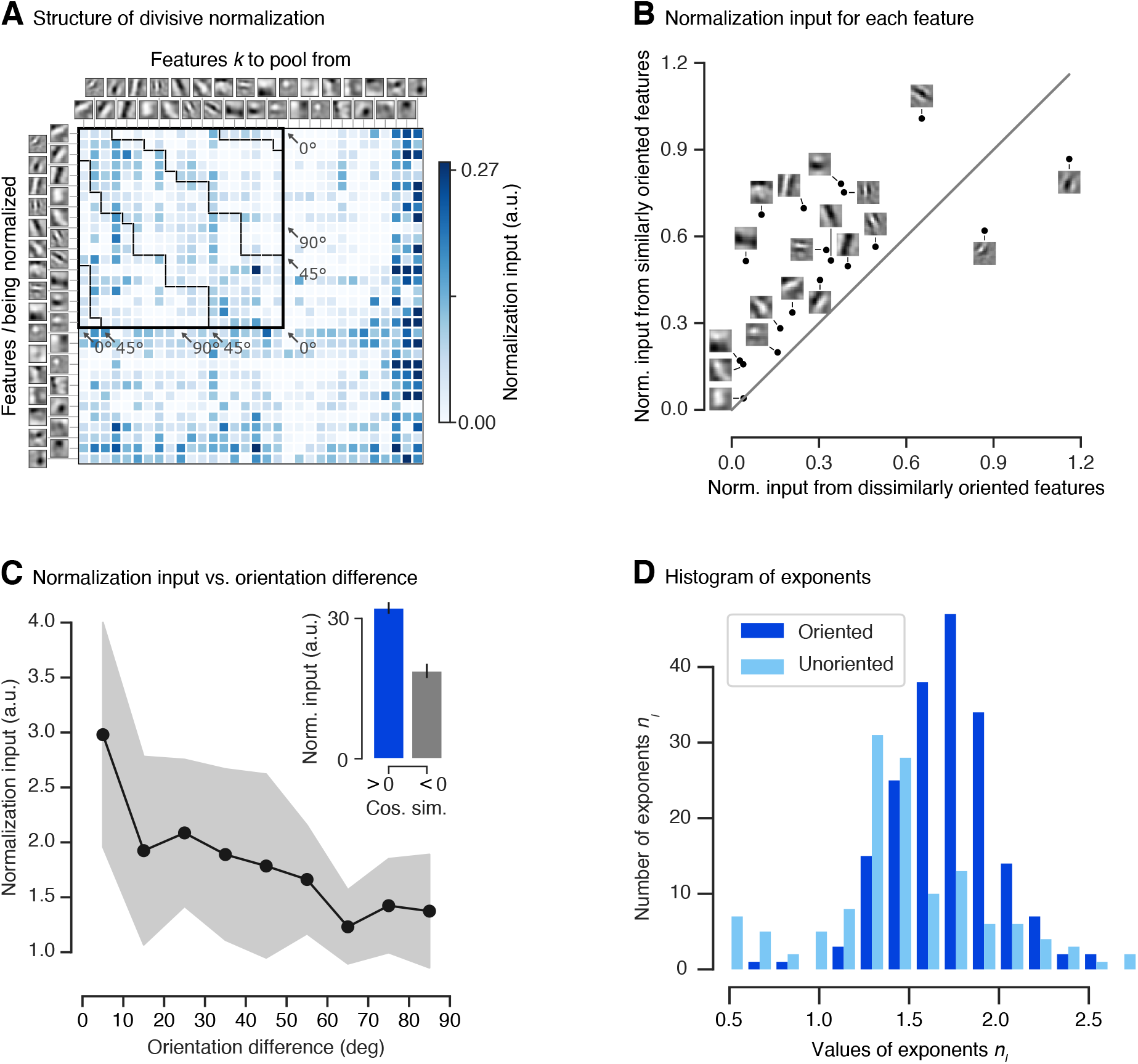
Structure of divisive normalization. **A**. The matrix shows the average strength of the normalizing inputs (products 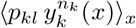 in denominator of Eq. (1) averaged across images; see Methods) for each combination of filter response being normalized (rows) and filter response providing normalizing input (columns). Darker shades of blue indicate stronger normalization. Orientation-selective filters are grouped at the top, ordered by preferred orientation and marked by the black square. The dashed black lines within the square separate pairs of filters with similar (< 45°) and dissimilar (> 45°) orientations. Normalizing inputs are stronger for similarly tuned filters. Unoriented filters mainly accounting for orientation-unspecific contrast are sorted by total normalization input. Darkest blue color corresponds to the maximum normalization input for the group of oriented filters, higher normalization input values for the unoriented filters are clipped to that value. Data of the model with highest accuracy on the validation set is shown. **B**. Normalization input from similar orientations (< 45°) compared to the normalization input from dissimilar orientations (≥ 45°) for each oriented linear filter. Grey line: identity. Most features are normalized preferentially by the responses of filters with similar preferred orientations. Data of the model with highest accuracy on the validation set is shown. **C**. Normalization input, binned into orientation difference of 10°. Each bin was averaged over the top-10 models (assessed on the validation set). The shaded area depicts the standard deviation per bin. **C inset**. Normalization input (norm. input) vs. cosine similarity between linear filters (cos. sim.) averaged across the top-10 models (assessed on the validation set). A cosine similarity greater than zero corresponds to similar features. Error bars: standard error of the mean. **D**. Histogram of DN exponents *n*_*l*_ (Eq. 1) of the ten best performing models in terms of validation set accuracy. Darker/lighter color: exponents corresponding to driving inputs due to oriented/unoriented linear filters. Most values are larger than one, with a few exceptions mainly corresponding to unoriented filters.

First, we coarsely quantified the difference with which these two groups contribute to normalization. We split the sum in Eq. (1) into two parts and collected the normalizing input of features with similar orientation as the driving feature (< 45°) and that of features with dissimilar orientations (≥ 45°). By analyzing the normalization of each oriented feature individually, we found that most oriented features are more strongly normalized by features with similar orientations (Fig. 5B). To assess whether our qualitative observation above is a general property of the data or a spurious characteristic of the best model chosen, we repeated this analysis for the top-10 performing models (assessed in terms of accuracy on the validation set, only differing in the initialization of the model parameters before training) and observed a similar behavior. Summing up the normalizing input across the features, we found that, for all of these models, similar orientations contributed more strongly than dissimilar orientations. Taking the data of all top-10 models into account, we found that similarly oriented features contributed (47 ± 10) % more normalizing input than dissimilar features (mean and 95% confidence interval over the ten best models selected in terms of validation set accuracy; Wilcoxon signed rank test: *p* < 0.006, *N* = 10 models; Cohen’s *d* = 1.2).

We further quantified the orientation-specific nature of normalization at a more fine-grained level. Instead of lumping orientation differences into two groups as before, we split up the normalizing inputs into nine bins of 10° width each and averaged those bins across the top-10 models. This analysis revealed that the strength of the normalizing inputs decreased as the difference in orientation increased (Fig. 5C). Hence, the more similar a normalizing feature’s orientation was to the feature to be normalized, the stronger was its contribution to normalization (linear regression analysis on all normalization input vs. orientation difference pairs of best model in terms of validation accuracy: *p* < 10^−6^). In fact, features in the group most similar to the driving input contributed (126 ± 35) % more than those in the orthogonal group (mean and 95% confidence interval over the ten best models selected in terms of validation set accuracy; Wilcoxon signed rank test: *p* < 0.006, *N* = 10 models; Cohen’s *d* = 1.9).

An immediate question that follows from our previous findings relates to the contribution of more general feature properties beyond orientation selectivity. Is the normalization input higher only for similar orientation, or is normalization input generally higher for similar filters, without pre-specifying a certain property like orientation? To answer that question, we split the input to normalization in terms of cosine similarity between all oriented and unoriented filters into a group of similar (cosine similarity > 0) and dissimilar (cosine similarity < 0) features. Summing up the normalizing input across the features and averaging across the best ten models, we found that in general similar features contributed (85 ± 47) % more normalizing input than dissimilar features (Fig. 5C inset; mean and 95% confidence interval over the ten best models selected in terms of validation set accuracy; Wilcoxon signed rank test: *p* < 0.006, *N* = 10 models; Cohen’s *d* = 2.9).

Analysing the exponents of our divisive normalization model, we found a mean value of 1.61 (averaged across the best ten models on the validation set) with most values being greater than one (Fig. 5D), which previously has been connected to an intensified winner-take-all behavior of responses (Busse et al., 2009; Carandini and Heeger, 2012). In a few exceptions exponents learned by our model are smaller than one, mainly corresponding to driving inputs from unoriented filters.

#### 2.4.1 Control: Nonspecific divisive normalization reduces accuracy

To determine how important orientation-specific normalization is, we performed a control experiment: For each feature *l* being normalized, we constrained all of its incoming normalization weights *p*_*kl*_ to be identical. This model is an instantiation of the previous models (Heeger, 1992; Carandini et al., 1997; Busse et al., 2009) assuming non-specific normalization from all features. Note that, since the feature orientations were non-uniformly distributed (Fig. 5A), this model’s normalization was weakly orientation tuned despite equal weights for all features (orientation difference < 45° contributed 14% stronger than ≥ 45°; Wilcoxon signed rank test; *p* < 0.007, *N* = 10 models; Cohen’s *d* = 0.3). This model achieved a performance of (41 ± 2) % between the baseline and gold standard ((48.9 ± 0.1) % FEV, mean and 95% confidence interval over the ten best models selected in terms of validation set accuracy). The unspecific DN model was included in the hypothesis test showing statistically significant performance differences between all model types (Section 2.2; pairwise Wilcoxon signed rank test on best models in terms of validation accuracy: *p* < 0.024, *N* = 166 neurons, family-wise error rate *α* = 0.05 using Holm-Bonferroni correction). While the unspecific control model does not match the performance of our more general DN model, it does outperform the subunit baseline. Thus, stronger orientation-specific normalization is necessary to achieve the full DN model performance, but a simpler form of uniform divisive normalization can account for a large fraction of the improvement over the subunit baseline.

#### 2.4.2 Control: All channels contribute to our model’s prediction

One potential caveat of our analyses so far is that we analyzed the orientation specificity of DN in terms of the convolutional feature maps in our model’s core rather than the actual neurons we recorded. These features provide a much more compact view of the population of neurons, because they are invariant to the receptive field locations and the neural responses are simple linear combinations of those features. Moreover, the model’s core offers a direct interpretation of its normalization weights. However, it is not clear a-priori whether all features are equally important for predicting the activity of the neurons in our population. Thus, considering convolutional features instead of actual neurons may lead to a skewed view of the population. To verify that this is not an issue, we quantified how much each feature contributed to the overall activity of all neurons by normalizing the feature readout weights across channels and averaging across neurons. The resulting distribution (Fig. 6) containing these averaged feature readout weights for the best ten models had a coefficient of variation of 0.2. We therefore concluded that all features were read out by roughly the same number of neurons and hence were similarly important to predict neural activity. Thus, our interpretation of orientation-specific normalization is unlikely to be an artifact of analyzing the convolutional features rather than the actual neurons.

**Figure 6:**
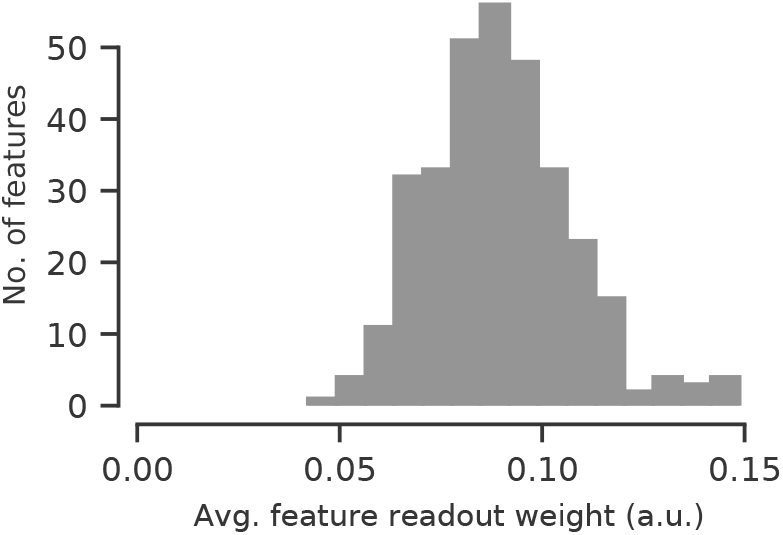
Histogram of feature readout weights of the ten best performing models in terms of validation set accuracy. For each model, feature weights are normalized across channels and averaged across individual neurons. All model’s channels are used to predict neural activity.

#### 2.4.3 Control: Surround interactions are unlikely driving orientation-specific normalization in the RF center

We observed orientation-specific divisive normalization in the receptive field. Surround suppression is known to be orientation-specific as well (Blakemore and Tobin, 1972; DeAngelis et al., 1994; Cavanaugh et al., 2002a,b; Coen-Cagli et al., 2015), so a potential concern would be that some of the extra-classical surround of a unit’s RF contributed to the results presented above. To rule out this possibility, we explicitly verified *in silico* that our model learning DN inside the RF learned no significant surround-suppression. For each neuron, we generated a circular grating stimulus with optimal orientation and spatial frequency, which we presented to our model. As we increased the grating diameters, our DN model’s spike count predictions were suppressed only for very few cells for very large stimuli (Fig. 7A, inset), suggesting only weak surround-suppression in very few cells. To quantify this observation, we computed a surround suppression index reporting the reduction from the highest measured response to the response for a grating completely filling the whole image for each neuron (Fig. 7A, inset). Thus, no suppression is indicated by a suppression index of zero, whereas a value of one would mean maximum suppression (see Methods for details; Cavanaugh et al. 2002a). Across the best ten DN models and all neurons, we found only very few exceptions from no suppression (Fig. 7A) with an average suppression index of (1.6 ± 0.8) % (mean and 95% confidence interval over the ten best models selected in terms of validation set accuracy). If our DN model would have learned surround suppression, we would have expected that the predictions of a substantial number of neurons would significantly decrease for larger grating sizes leading to substantially higher suppression indices (Cavanaugh et al., 2002a), which is not the case for our DN model. In conclusion, we expect the influence of very few neurons showing mostly very weak surround-suppression to our results to be very small making it unlikely that surround suppression could have led to the substantial orientation-specific normalization we observed. For the non-specific DN model and the three-layer CNN we found very similar behavior.

**Figure 7:**
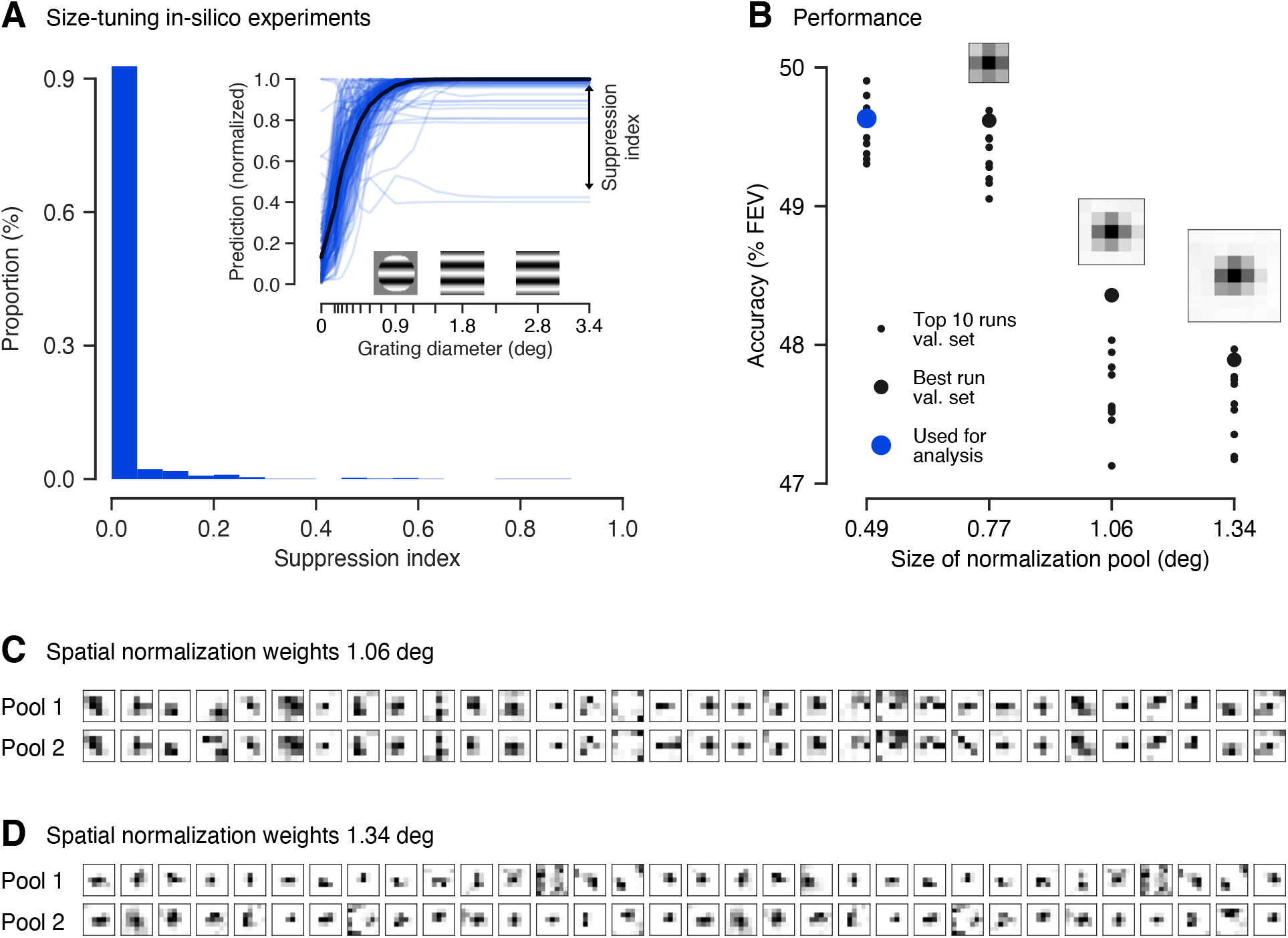
**A**. Size-tuning *in silico* experiments. **A. inset:** Prediction of the best DN model (chosen by validation set accuracy) for all neurons to gratings of increasing size. The gratings’ properties were determined from the units’ optimally stimulating Gabor pattern. As grating diameter increased, only very few neurons showed mostly weak suppression. Predictions normalized to maximum response per neuron. Suppression index measures asymptotic suppression relative to the maximum prediction **A. main panel:** Across all neurons and the ten best DN models (chosen by validation set accuracy), almost no neurons show significant surround suppression. **B**. Test set performance of the ten best performing DN models. The model’s performance rapidly decreases for spatially increasing normalization pool size (in units of visual angle in degrees). The best model on the validation set is indicated by a blue dot. **C. & D**. Weights of the spatial normalization pool for the best performing model with pool size of (**B**.) 1.06°of visual field (5 px × 5 px) and (**C**.) 1.34°of visual field (7 px × 7 px; all evaluated in terms of the validation set accuracy). For each feature (columns), the two components (rows) of the in total 32 spatial normalization pools are shown. Darker color corresponds to higher weights. Both components are similar. **B. insets:** Average across features and normalization pool components. The model learned normalization from the receptive field center (on average).

As a further control to verify that the orientation-specific normalization we observed in the RF is unlikely to be caused by surround influence, we fit a more general DN model which could additionally learn the spatial structure of the normalization pool, instead of limiting it only to neurons with overlapping receptive fields. To account for the RF’s surround, we spatially expanded the model’s normalization weights *p*_*kl*_ → *p*_*kluv*_ and introduced a convolution across space into the normalizing sum in Eq. (1):

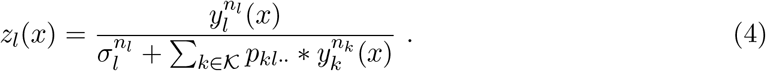

Here, the normalization weights *p*_*kl*··_ = *p*_*kluv*_ encode which input drives 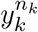 are pooled over what spatial extent (indexed by *u, v*) to normalize the input drive 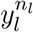 .Note that for the special choice of normalization weights *p*_*kl*11_ we perform a *u* × *v* = 1 × 1 convolution resulting in Eq. (1) describing a DN model without surround. To keep the number of parameters computationally tractable, we factorized the normalization weights *p*_*kluv*_ allowing for two normalization pools (indexed by *m*) per output channel *l*, each consisting of a set of spatial integration weights *c* and feature weightings *d*,

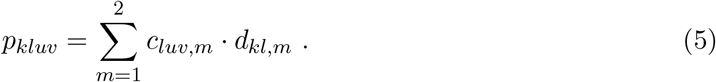

We constrained *c* and *d* to be non-negative to make sure the denominator in Eq. (4) is non-negative (recall that 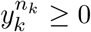 due to Eq. (2)). The two normalization pools indexed by *m* could have different patterns of weights along the feature and spatial dimensions. Our extended model is therefore general enough to account for the standard model of DN with a nonspecific center normalization pool and orientation-specific surround suppression (see Methods for details).

In line with what we expect from our small stimuli mostly covering the RF center, spatially expanding the normalization pool to cover larger surround areas did not increase our model’s accuracy; in fact, for lager normalization pools the performance even decreased (Fig. 7B; for the best models on the validation set statistically significant beyond 0.77° of visual field, pairwise Wilcoxon signed rank test: *p* < 0.025, *N* = 166 neurons, family-wise error rate *α* = 0.05 using Holm-Bonferroni correction). This could be due to the increased number of parameters, which might lead to over-fitting or non-optimal learning, since adding more weights to the model makes optimization harder. The best performance was achieved by models with the smallest normalization pool mostly within the units’ RFs (0.25° – 0.75° diameter estimated by spike triggered average, see Methods; 0.86° estimated from the smallest grating that leads to at least 95% of the maximum response in our *in silico* analysis, averaged across neurons and the best ten DN models with smallest normalization pool). The normalization weights of the extended spatial normalization pools showed no visible separation into center and surround and exhibited no or only weak contributions from the RF’s surround (Fig. 7). If leakage from the surround was the source of orientation-specific normalization, we would expect to observe spatial profiles that cover the surround, which is not the case (except for a few exceptions, which are mostly untuned features).

In summary, we controlled that the orientation-specific normalization learned by our spatially constrained DN model is unlikely caused by orientation-specific influence from the RF surround but mainly from the RF center. The reason for this limitation could be in our dataset, as the stimuli we used were mostly restricted to the receptive field masking out any surround influences and their short presentation time compared to the delay after which surround influences kick in (see Discussion and Methods), preventing any model to capture information from the surround.

## 3 Discussion

To improve our understanding of primary visual cortex, we asked what condensed nonlinear function state-of-the-art CNNs might implement for predicting V1 responses to localized natural stimuli. To answer this question, we developed an end-to-end learnable divisive normalization model and fit it to neural responses. Both the unspecific control model and the full model that learned the normalization pool outperformed the subunit model, with the full DN model outperforming its non-specific variant. Through the superior performance of the DN model, we quantify the relevance of divisive normalization for predicting V1 responses to natural images. With the help of *in silico* experiments, we found that both, the DN and the CNN model learned cross-orientation inhibition, showing that the deeper layers of the CNN model approximate the DN nonlinearities. Compared to the multi-layer CNN, the DN model offers a direct, easily understandable interpretation of its normalization pool. For the DN model, we found that normalization by similar orientations, and in general by similar features, is higher compared to dissimilar features within the receptive field.

Along those lines, one may ask whether the difference between the non-specific DN model and the full model that learns orientation specific normalization weights is relevant, because the full model may simply be able to better account for some insignificant biological heterogeneous imperfection due to its additional parameters. We believe that this explanation is unlikely, because oriented features are preferentially normalized by channels with similar orientation. If the model was simply picking up some biological imperfection, we would expect random deviations from uniform normalization weights rather than weights that depend systematically on preferred orientation.

Previous experimental work investigated suppressive phenomena within the receptive field only qualitatively or used simple stimuli that mainly consisted of a combination of driving and mask gratings. Morrone et al. (1982) found suppression at all orientations, but did not investigate orientations similar to preferred orientation. Bonds (1989) reported predominantly orientation-nonspecific suppression, although three of fourteen cells exhibit stronger suppression with masks oriented similarly to the neurons’ preferred orientations, and a few other cells were suppressed most strongly by mask orientations orthogonal to the preferred orientation. Similarly, DeAngelis et al. (1992) found suppression to be predominantly independent of orientation, although for some cells an increased suppression for a range of orientations near the optimal excitatory orientation was apparent. Heeger (1992) explained those results by proposing an orientation-nonspecific divisive normalization model. Carandini et al. (1997) considered the possibility of orientation-specific normalization which provides a marginal improvement in the quality of their model fits to the data. However, they concluded that their dataset was not specifically designed to provide a strong test of this question and their results were inconclusive in this respect. Busse et al. (2009) developed a quantitative model for the response of a population of neurons to a combination of gratings. Assuming nonspecific normalization by overall contrast, their model predicted the collective action of the whole neuron population better than linear and winner-take-all baselines, but they did not test against an orientation-specific alternative model. To summarize, these studies found phenomena that are predominantly explained by nonspecific normalization (Heeger, 1992), some of them encountered only weak orientation-specific phenomena and only in relatively few cells. Until now, a quantitative analysis of orientation-specificity on a data set of natural images was missing.

Our findings are largely consistent with previous experimental results and quantitatively refine them using a larger dataset, place them in the context of other models of V1 and show that the same normalization mechanisms observed with simple stimuli also apply under more natural stimulus conditions. Interestingly, and somewhat in contrast to earlier work, features with preferred orientations within 10° of the driving feature provided twice as strong normalizing input than those with orthogonal preferred orientations.

The reason for this difference between our findings and previous studies could be that we used natural stimuli, which have different image statistics compared to simple stimuli used in earlier studies. Furthermore, most previous studies of divisive normalization were performed in cats (Morrone et al., 1982; Bonds, 1989; DeAngelis et al., 1992; Busse et al., 2009) and the results therein may not generalize to monkeys, for which preceding studies are inconclusive regarding orientation specificity (Carandini et al., 1997).

Recent work modeling a large set of classical psychophysical data also suggests an orientation-specific divisive normalization: Schütt and Wichmann (2017) developed an image-computable model of early vision very similar in structure to ours, and found that in order to explain classical data on contrast detection, contrast discrimination and oblique masking, their model required divisive normalization to be orientation-specific. Similar results had been reported in an earlier study (Itti et al., 2000). In contrast to those psychophysical studies involving experiments with human observers, in our paper we studied individual spiking neurons in primary visual cortex in response to natural stimulation, providing physiological evidence for orientation-specific divisive normalization. Moreover, the previous psychophysical models used a predefined Gabor filter bank and assumed that divisive normalization’s orientation specificity is Gaussian distributed, learning only the standard deviation of the distribution. In contrast, our model is far more flexible, not making such strong assumptions, relying solely on the data to learn oriented filters and their divisive interactions.

Following a normative approach, Schwartz and Simoncelli (2001) derive an ecologically justified divisive normalization model from the efficient coding hypothesis (Barlow, 1961) that is able to qualitatively describe the orientation masking data of Bonds (1989). Reducing the statistical redundancy of responses to natural stimuli predicts that normalization should be stronger for neurons that exhibit a higher dependency in their unnormalized responses. This theoretical result implies that normalization weights should not be uniform, consistent with our empirical findings. Iyer and Burge (2019) compare the output statistics of a divisive normalization model with broadband and narrowband normalization to natural images, 1/*f* stimuli and white noise stimuli. They report that feature-specific divisive normalization improves the discriminability among natural images compared to unspecific normalization.

Is our discovery of divisive normalization by similar orientations actually implemented by the connectivity of neurons in primary visual cortex? The answer to this question could be reflected in the connectivity from inhibitory parvalbumin-expressing (PV) interneurons to pyramidal cells and their relation to neurons’ tuning properties. Hofer et al. (2011) find that, in the mouse, pyramidal cells and PV cells are homogeneously connected. Although a weak bias towards orientation tuning is apparent, they conclude that local inhibition in V1 is primarily non-specific. However, despite the connection probability between PV and pyramidal cells being homogeneous, connection strengths are quite heterogeneous: Individual PV cells strongly inhibit those pyramidal cells that share their visual selectivity (Znamenskiy et al., 2018). This result is in line with our finding of orientation-specific normalization. Similarly, recent work employs single-neuron perturbations in layer 2/3 of mouse V1 and directly measures higher suppression of neighbouring excitatory neurons with a similar preferred orientation (Chettih and Harvey, 2019).

The stimuli in our dataset were specifically designed to investigate nonlinear processing in the RF center and to minimize surround suppression. The control analyses showing that surround suppression is unlikely in our dataset serve only the purpose of verifying that our conclusions about orientation-specific normalization in the receptive field are not due to leakage of surround suppression (which is known to be orientation-specific; Blakemore and Tobin 1972; DeAngelis et al. 1994; Cavanaugh et al. 2002a,b; Coen-Cagli et al. 2015). Without question, surround suppression is an important component of nonlinear processing in V1 and deserves further investigation and modeling using a dataset designed for that purpose.

In our stimuli the fully visible part was spatially restricted to approximately the size of the receptive field. For our dataset the RF diameters of 0.25° – 0.75° have been roughly estimated by spike-triggered average. RF size estimates based on the grating summation field (GSF) tend to be larger by approximately a factor of two Cavanaugh et al. (2002a). The GSF is defined by the smallest grating diameter that leads a unit to respond with at least 95% of its maximum activity (Cavanaugh et al., 2002a). Accordingly, the average GSF size we estimated based on our *in silico* size-tuning experiments had a diameter of 0.86°, roughly corresponding to the fully visible part of our stimuli (1° of visual field). For the remaining stimulus portion (1° – 2°) outside of the GSF there could be normalizing influences from the surround (Cavanaugh et al. 2002a Fig. 1A and Fig. 2). However, it is rather unlikely that this region strongly influenced our data, as in our stimuli a cosine mask fading to zero was applied in this area (see Methods). Rather, we would expect significant surround effects beyond 2° of visual field, which is not covered by our stimuli. Furthermore, in the results we have shown here, our model saw only the central 1.14° of the images (except for the variants extended to the surround, which did not learn significant surround interactions either). Thus, as we expected, our DN model did not exhibit significant surround suppression in our size-tuning control experiments and spatially extended DN control models suggested no surround interaction (Fig. 7). Summarizing our control analysis, literature suggests that in general there are surround interactions under stimulation with natural images (Coen-Cagli et al., 2015; Vinje and Gallant, 2000), however, our spatially constrained stimuli likely restricted us to study DN within the receptive field and very likely prevented us from learning the influence of the RF’s surround on normalization, making the surround unlikely to affect our result of orientation-specific normalization within the RF center.

Another property of our dataset that minimized the effect of surround suppression is that we focused on single images to predict a spike count in a relatively short time window covering the transient response, and ignored any temporal aspects or more sustained periods of the response. If there was some interaction from the partly masked surround, we would expect the onset of these effects to be delayed. Several studies report that the onset of surround suppression exhibits a 15 – 60 ms delay relative to the onset of center RF responses (Angelucci and Bressloff, 2006). Bair et al. (2003) reported average delays as short as 9 ms, however, its peak effect was typically reached 40 – 50 ms after its onset. As responses to the center RF occur on average 52 ms after stimulus onset (Bair et al., 2003), in the dataset we used (Cadena et al., 2019) we would expect significant surround influences to set in at the end of the recording window, which was 40 – 100 ms after stimulus onset. Hence, if there might be any contribution from the surround, we would expect it to contribute only very weakly to the overall spike counts determined in this time window. Note, the limitations in studying surround effects are imposed by the available data – the modeling approach may well generalize to cover both the surround and the temporal structure – and thus the extra-classical receptive field should be addressed in future work with a dataset containing larger stimuli.

Compared to our divisive normalization model, the three-layer CNN performs a few percentage points FEV better, showing that it captures additional (non-linear) phenomena that divisive normalization cannot account for. Future work will be necessary to find out what these unexplained effects are. This result shows the importance of quantitative, data-driven modeling: our work suggests that a model that can account for pooling of subunits (complex cells) and divisive normalization is structurally insufficient to achieve optimal performance in modeling V1 responses.

The performance difference between subunit and DN model measured in terms of FEV is also just a few percentage points. However, the phenomenological difference between those models is quite substantial, as the DN model correctly captures cross-orientation inhibition while the subunit model does not. Therefore, one should probably not only measure model performance by considering FEV over a set of randomly sampled natural images, but instead also consider other metrics or specifically chosen test sets that highlight certain aspects of neural computation – such as cross-orientation inhibition.

Important future research directions that follow our methods include studying the role of divisive normalization in higher areas of the ventral stream, like V4 and inferotemporal cortex. Building a predictive model by stacking successive DN layers or including a final divisive nonlinearity for the shared feature space are great candidates for future consideration, especially as earlier studies already report signs of normalization in these higher areas (Zoccolan et al., 2005; Kaliukhovich and Vogels, 2016; Reynolds and Desimone, 2003).

In conclusion, we developed a model consisting of one layer of subunits followed by learned orientation-specific divisive normalization, which accounted remarkably well for V1 data. We hope that this quantitative approach of evaluating theories of computation in the brain by formalizing them as (components of) trainable predictive models will be used more widely in the future, so the field will (slowly) converge to an accurate general-purpose model of the visual system applicable to natural inputs.

## 4 Methods

### 4.1 Experimental details

We used the dataset described in detail in Cadena et al. (2019) and provide a summary of the most important characteristics here. Electrophysiological recordings from two healthy, male rhesus macaque monkeys aged 12 and 9 years were performed with a 32-channel linear silicon probe. The monkeys were head-fixed and placed in front of a screen. They were trained to fixate on a target located at the center of the screen. The start of a trial was determined by maintained fixation on the target for 300 ms. The fixation tolerance was set to 0.42° around the center of the target. At the beginning of each recording session, population receptive fields were mapped with a sparse random dot stimulus. Each dot was of size 0.12° of visual angle and was presented over a uniform gray background, changing location and light intensity (black or white) randomly every 30 ms. The receptive field profiles per electrode channel were then obtained via reverse correlation (i. e. spike-triggered average). The center location of the population receptive field was subsequently estimated by averaging over channels and fitting a two-dimensional Gaussian to the reverse correlation profiles. Afterwards, this location was used to place the images of the natural stimulus paradigm.

The dataset in Cadena et al. (2019) consists of 7 250 distinct natural, greyscale images which were presented two to four times each. A fifth of these images (1 450) were taken from ImageNet (Russakovsky et al., 2015). Four additional texturized images were synthesized from each of them, preserving varying degrees of higher-order statistics. The images were cropped to 2 × 2 degrees of visual angle (140 px × 140 px). Before displaying the images on the screen, the images were normalized such that the central 1° (70 px) of each image had the same mean (111.5) and standard deviation (45) determined across the central portion of all original images. Pixels with an intensity that fell outside the display’s range [0, 255] where clipped. Afterwards, all images were overlaid with a circular mask with a soft cosine fade-out fading to the screen’s mean gray intensity (128) and an aperture with a diameter of 1°.

Images were presented for 60 ms with no blanks in between. Neural responses were extracted in time windows of 40–100 ms after image onset (Fig. 2), accounting for typical response latencies in primary visual cortex. The image sequence was randomized with the restriction that consecutive images do not belong to the same type (i. e. natural or one of the four texturized versions).

A few isolated neurons were discarded if their stimulus driven variability was too low Cadena et al. (2019). The explainable variance in a dataset is smaller than the total variance because the observation noise prevents even a perfect model to account for all the variance in the data. Thus, targeting neurons that have sufficient explainable variance is necessary to train meaningful models of visually driven responses. For a neuron’s spike count *r*, the explainable variance Var_exp_[*r*] is the difference between the the total variance Var[*r*] and the variance of the observational noise 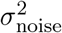,

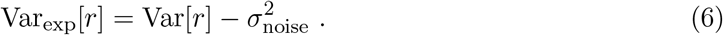

We estimated the variance of the observational noise by computing the variance of a neuron’s response *r*_*t*_ in multiple trials *t* in which we presented the same stimulus *x*_*j*_ and subsequently taking the expectation *E*_*j*_ over all images,

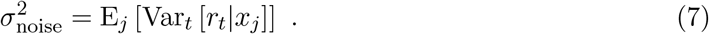

Neurons for which the ratio between the explainable to total variance was below 0.15 were removed. The resulting dataset includes spike count data for 166 isolated neurons, with an average ratio of explainable to total variance of 0.285. These neurons were recorded at 1° − 3° eccentricities and estimated receptive field size diameters were between 0.25° and 0.75°. Since RF sizes were roughly estimated using spike-triggered average, it is likely that the values reported here underestimate the grating summation field defined by the smallest grating diameter that leads a unit to respond with at least 95% of its maximum activity by a factor of approximately two (see Discussion; similar to the minimum response field (MRF) underestimating the GSF as reported by Cavanaugh et al. 2002a).

To keep the results of our models consistent and comparable to the gold standard baseline from Cadena et al. (2019), we down-sampled the images by a factor of two to train our models. Likewise, images were cropped symmetrically, keeping the 40 × 40 central pixels (1.14° of visual angle). This size covers all of the recorded neurons’ receptive fields, with a slight variability in their spatial location. Furthermore, the stimuli light intensities across all pixels and all images were centered around zero and normalized to have unit standard deviation. Additionally, we used the same random dataset splits of Cadena et al. (2019) into training (64%), validation (16%) and testing (20%). We assessed our models’ accuracy for a specific architecture or set of hyper-parameters in the validation set and we report performance on the test set. We consistently used the same split throughout our study.

### 4.2 Divisive normalization model

Our model consists of two parts, a nonlinear core and a linear readout (Section 2.1 and Fig. 1). The core (Fig. 1A) processes the input stimulus *x* by convolving it with 32 filters *w*_*k*_ of size 13 px × 13 px (0.37° of visual angle) without padding, defining a bank of features indexed by *k*. Subsequently, we apply batch normalization (Ioffe and Szegedy, 2015) without re-scaling (BN*) which adds a bias term (*o*_*l*_ in Eq. 2) and scales the responses to be of unit variance. This operation does not affect the overall computation and biological interpretation of our model, since scaling the driving inputs (*y* in Eq. 1) by a factor *β* whilst scaling the normalization weights (*p* in Eq. 1) and linear readout weights (Eq. 3) by 1*/β* yields the same output. The batch normalization step is followed by a rectified linear unit (ReLU) nonlinearity

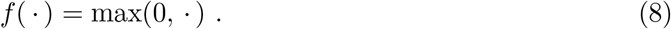

Hence, equivalent to Eq. (2), the resulting 32 feature maps of size 28 px × 28 px for the excitatory drive are given by

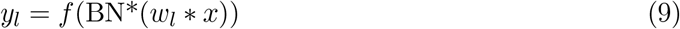

and contain information about the full input space covering 1.14° of visual field. Many neurons perform similar computations but respond at different localized areas of the visual field. Those receptive fields are represented by the kernels *w*_*l*_, which we implemented convolutionally to make use of this knowledge. Furthermore, the ReLU nonlinearity (Eq. 8) ensures that all feature maps are non-negative, *y*_*l*_ ≥ 0 being coherent to the biological interpretation of an excitatory drive.

The feature maps *y*_*l*_ are then normalized divisively to produce 32 output feature maps

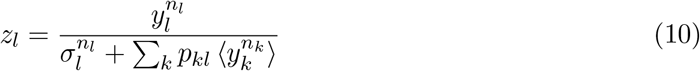

shared by all neurons. Here, all operations are element-wise and the scalar semi-saturation

constant *σ*_*l*_ ≥ 0 is learned from the data. To include normalization by other channels *k*, we first exponentiate the excitatory feature maps *y*_*k*_ by the scalar *n*_*k*_ ≥ 0 element-wise, which is learned from the data as well. Subsequently, low-pass filtering 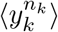 is performed through average pooling in space with pool-size 5 px × 5 px (0.14°, covering 0.49° in the input space taking the convolutional layer into account). We perform this pooling in order to achieve (approximate) phase invariance of the normalizing input without requiring a large number of filters with different phases. Subsequently, the results of the low-pass filtering are summed up, weighted by the normalization weights *p*_*kl*_, and added into the denominator, resembling Eq. (1). Furthermore, the normalization weights are constrained to be non-negative, *p*_*kl*_ ≥ 0. Together with *y*_*k*_ ≥ 0 and *σ*_*l*_ ≥ 0, this ensures that the denominator in Eq. (10) is non-negative, hence having a well-defined biological interpretation.

We converted the core’s output feature maps *z*_*l*_, shared by all neurons, to the activity of individual neurons via a readout for each of them (Fig. 1B). To do so, we factorized the readout into spatial readout weights *a*_*uv,i*_ ≥ 0 and feature readout weights *b*_*l,i*_ ≥ 0 that pick the relevant locations and features plus a bias *q*_*i*_,

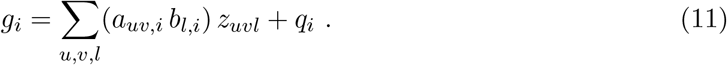

Here, *u, v* index space and *i* indexes neurons. This factorization is beneficial because it reduces the number of parameters in the readout. Also, we wanted to ensure that the readout does not model any complex computations, which we achieved by this factorization and the non-negativity of the readout weights. Additionally, we limited complexity by imposing a sparseness prior on both weights, because each neuron should only respond to its receptive field which is represented by a sparse spatial readout weight and should not mix many different features which corresponds to a sparse feature readout weight. The readout can, however, model a complex cell (Hubel and Wiesel, 1962) by linearly combining multiple channels of the shared feature space.

We fitted an output nonlinearity to obtain the final prediction of a neuron’s activity 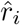 as suggested by Cadena et al. (2019) from which we paraphrase the description here. Optimizing the output nonlinearity improves data-driven models, but has to be done carefully when simultaneously learning the shared core and the readout weights end-to-end. We therefore construct the output nonlinearity inspired by the shifted exponential linear unit (ELU*; Clevert et al. 2015),

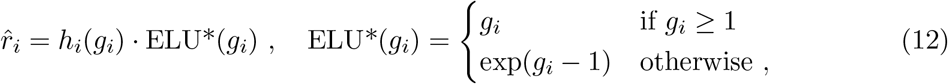

keeping the resulting activity non-negative, and multiply it by a non-negative function

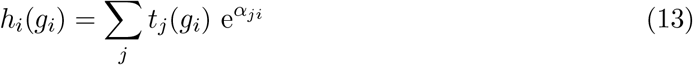

with parameters *α*_*ji*_ which are learned for each neuron *i* independently. The basis functions *t*_*j*_(*x*) in the argument of *h* are tent functions leading to a piecewise linear function inside the exponential. The tent functions are defined as

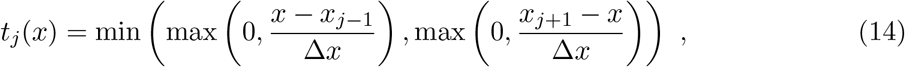

with interpolation points *x*_*j*_ = −3, −2.82, …, 6 and Δ*x* = 0.18. By regularizing the coefficients *α*_*j*_ with a *L*_2_ and a smoothness penalty, the model is pushed towards using the identity function for inputs larger than one and a shifted exponential function for smaller inputs. If the data provides strong evidence in favor of a different output nonlinearity, the piecewise function *h*_*i*_ can be used to modify it. For example, the output nonlinearity is general enough to learn a sigmoidal shape, although for our best-fit models across model types it learned a slight expansive deviation from the identity in most cases and in a few cases a mix of slight expanding and slight compressing deviations from the identity function.

To optimize our model’s parameters, we maximized the log-likelihood of the model’s predictions given the data. To do so, we assumed that neurons’ spikes are produced by a Poisson process. Our model predicts the average spike count 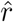 of a neuron, hence the probability of observing *r* spikes in the experiment is

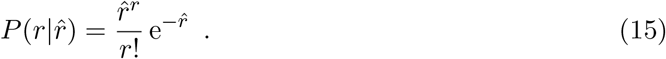

From that follows the Poisson log-likelihood

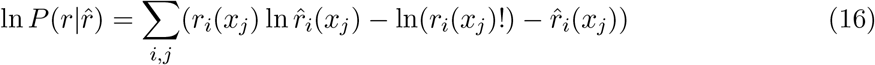

for all neurons *i* and all stimuli *x*_*j*_. A neuron’s response *r*_*i*_ ≡ *r*_*i*_(*x*_*j*_) depends on the stimulus *x*_*j*_, which we suppress in our further notation for better readability. For implementation reasons, we wanted to minimize the Poisson loss function

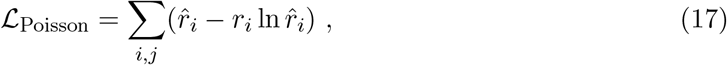

which is the negative of the Poisson log-likelihood (Eq. 16), where we omitted ln(*r*_*i*_!) since this term does not depend on our model.

Furthermore, three terms regularizing the model’s parameters were applied to the loss. We imposed a smoothness prior on the kernels *w*_*k*_ to ensure the spatial continuity of the predictors’ receptive fields. The according penalty on the loss for weights being not smooth was determined with a Laplace filter *L* to be

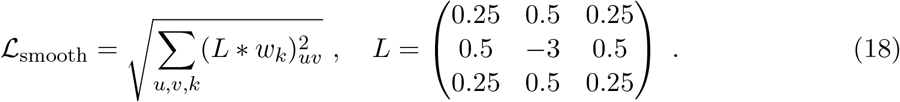

Due to their receptive fields, neurons only respond to a small, localized area of the visual field, which is why we imposed a sparsity prior on the spatial readout weights *a*_*uv*_. Furthermore, neurons should only pool from a small set of feature maps to ensure that the readout does not perform complex computations. Thus we imposed a sparsity prior on the feature readout weights *b*_*l*_ as well. We achieved this by adding the *L*_1_-norm of both weights

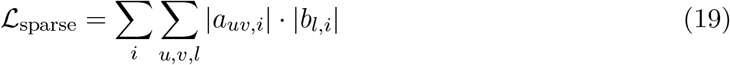

to the loss function. As described by Cadena et al. (2019), we regularized the output nonlinearity for all neurons by penalizing the sum of squares of the first and second finite differences of the weights *α*_*ji*_ to keep the learnable exponential function *h*_*i*_(*x*) smooth and close to 1,

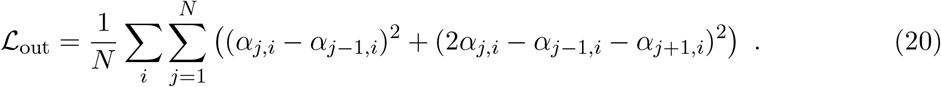

The final loss function to minimize with respect to our model’s parameters is

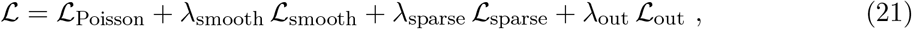

where *λ*_smooth_, *λ*_sparse_ and *λ*_out_ are hyper-parameters which set the strength of the smoothness, the sparsity and the output nonlinearity penalty, respectively.

### 4.3 Divisive normalization model extended to normalization from surround

To extend our DN model to capture normalization from the spatial surround of a unit’s RF, we replaced the weighted sum accounting for normalization (Eq. 10) by a convolution that also covers space, keeping the rest of the original DN model unchanged,

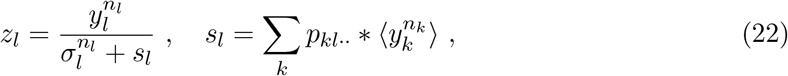

equivalent to Eq. (4). The new shared feature maps *z*_*l*_ consist of all element-wise operations where the newly introduced normalization feature maps *s*_*l*_ represent the strength by which the excitatory drive 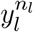 is normalized. The normalization feature maps are the result of a convolution between 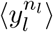 and normalization pool kernels *p*_*kluv*_.

For a larger convolutional kernel *p*, the feature maps *s* have smaller spatial dimensions than the excitatory feature maps *y* due to the convolution without padding. To be able to perform the element-wise division, we symmetrically cropped the excitatory feature maps *y* so that the resulting feature maps had the same spatial dimensions as *s*. Additionally, we wanted to keep the complexity (number of parameters) of the linear readout constant for all the size choices of the normalization kernel *p*. To this end, we symmetrically cropped the input images to a size that corresponds – after a forward pass through our model – to a shared feature space of spatial dimensions 28 px × 28 px (0.80°, covering 1.14° of the input image taking the convolutional layer into account). Overall, this process enabled a fair comparison across all sizes of *p*.

To keep the kernel size of *p* computationally tractable, we used dilated convolutions which have defined gaps between the weights of the convolutional kernels. We chose to skip four pixels between each weight which is reflected by a dilation factor of five. Thereby, we were able to pool from a relatively large extra-classical RF while using few parameters. If we would compute this convolution directly on the feature maps 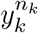, the dilation would cause us to skip some elements in the feature map 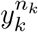 in the convolution’s inner product for one specific position of the convolutional kernel, i. e. for one specific element in the suppression feature maps *s*_*l*_. To consider all those elements we preceded the inner product computation of the convolution with an average pooling of 5 px × 5 px pool size (same as the dilation factor; 0.14°, covering 0.49° in the input space taking the convolutional layer into account) which we performed at every spatial position in the input feature maps 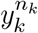. Then, exactly one weight of the convolutional kernel accounts for one 5 px × 5 px pool. In this view, the pools of the dilated kernel’s neighbouring weights have coinciding boundaries. So in addition to implementing shift invariance (see Section 4.2), the average pooling makes sure that we do not loose information for the extended DN model. Due to this pooling, a normalization kernel *p* of spatial size 3 px × 3 px would spatially cover a normalization pool of size 15 px × 15 px (0.43°, covering 0.77° in the input image taking the convolutional layer into account). We investigated models with normalization kernel sizes of 1 px × 1 px, 3 px × 3 px, 5 px × 5 px and 7 px × 7 px which spatially covered a five times larger normalization pool due to dilation. Those normalization pools covered visual angles of 0.49°, 0.77°, 1.06° and 1.34°, respectively.

### 4.4 Baseline models

#### 4.4.1 Convolutional neural network

Since the divisive normalization computation in our model was completely learned from the data, we wanted to compare to a baseline model that is purely data-driven as well. For this, the current state-of-the-art model is a convolutional neural network with three layers (Cadena et al., 2019). Its first convolutional layer consists of a kernel with spatial size of 13 px × 13 px (0.37° of visual field) and for the second and third layer of size 3 px × 3 px (0.09°) each, covering 0.43° and 0.49° of the input image, respectively. All layers use 32 channels, batch normalization (Ioffe and Szegedy, 2015) and ELU nonlinearity (Clevert et al., 2015),

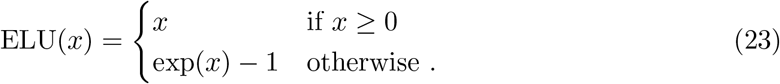

Similar to our model’s architecture, the core part of the CNN model results in a nonlinear feature space shared by all neurons which is mapped to each neuron’s activity with individual readout weights factorized in spatial and feature weightings. Sparseness of both of them is achieved by adding an *L*_1_-penalty to the according loss function. This readout employs the same final output nonlinearity but differs from ours in having no non-negativity constraints on the weights.

#### 4.4.2 Convolutional subunit model

Our convolutional subunit baseline model is structurally a one-layer convolutional neural network with multiple filters followed by a readout. It is exactly the same as our divisive normalization model (Section 4.2) but with the normalization function (Eq. 10) replaced by the identity function

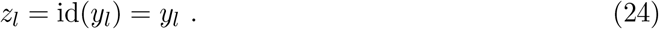

Hence, the only difference to our DN model is the lack of normalization. The shared feature maps *z*_*l*_ consist of rectified outputs of linear filters (Eq. 9) which approximate simple cells. The subsequent readout can sum up those simple cell responses with additional weightings, enabling the model to approximate complex cells (Hubel and Wiesel, 1962). We trained the model with the same loss function (Eq. 21) as the divisive normalization model.

### 4.5 Hyper-parameter optimization

Our model’s accuracy depends on several hyper-parameters. We set the initial learning-rate to 10^−3^ and used an early stopping training scheme: We evaluated the Poisson loss (Eq. 17) every 100 training steps, after ten iterations of no improvement we decayed the learning-rate by a factor of three, and we repeated this four times. For the filters *w*_*k*_ in the first convolution, we found that a size of 13 px × 13 px (0.37° of visual field) was optimal, the same is true for the number of 32 channels.

The weights *λ*_smooth_ of the smoothness penalty (Eq. 18), *λ*_sparse_ of the readout sparsity penalty (Eq. 19) and *λ*_out_ of the output nonlinearity’s penalty term (Eq. 20) in the full loss function (Eq. 21) were extensively cross-validated using the validation set of our data (Section 4.1). We randomly sampled the smooth-weight *λ*_smooth_ from a logarithmic uniform distribution in the interval [10^−9^, 10^−3.5^] and the readout sparse-weight *λ*_sparse_ from a logarithmic uniform distribution in the interval[10^−9^, 10^−4.^ for all models. For the divisive normalization model constrained to the receptive field center and the subunit model, we drew the output nonlinearity penalty weight *λ*_out_ from a logarithmic uniform distribution in the interval [10^−8^, 10^2^]. For those two models, we sampled 1 000 runs. For the non-specific divisive normalization model and the DN models extended to the spatial surround, we narrowed down the relevant parameter space of *λ*_out_ and sampled them from a logarithmic uniform distribution in the interval [10^−5^, 10^0^]. Since this halved the search space of the hyper-parameters, we halved the number of samples drawn for those models. Thus, the density of sample points in the search space was the same for all models and should lead to a fair comparison between their performance. For a fair comparison of all models in this study to the state-of-the-art three-layer CNN, we retrained this model with the same optimization procedure. After running optimization for a few 100 hyper-parameter samples, we observed that the best performing models were close to the upper bound of the readout sparsity penalty weight. Thus, we discarded those data-points and started searching again with a shifted search-interval for *λ*_sparse_ ∈[10^−5.5^, 10^−1^], sampling 1 000 model runs as before. In layer 2 and 3, we set the smooth-weights to zero and group-sparsity weights to 2.5 · 10^−4^ since Cadena et al. (2019) found best performance for those values and these parameters are specific to the three-layer CNN, being not applicable to the other models used in this study.

### 4.6 Accuracy evaluation

#### 4.6.1 Average correlation

For architecture search, hyper-parameter optimization and the selection of specific models for analysis we evaluated models’ accuracies on the validation set with the Pearson correlation coefficient between the measured spike counts and our models’ predictions, averaged over neurons. If the prediction for one neuron is constant, the according standard deviation is zero. Hence, for such neurons the correlation coefficient was not computable due to division by zero. For those neurons, we set the correlation coefficient to zero before averaging. This average correlation measure does not consider observational noise (Eq. 7).

#### 4.6.2 Fraction of explainable variance explained

For reporting accuracy values in this paper, we used the data’s test set to compute the fraction of explainable variance explained (FEV) per neuron

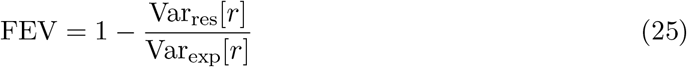

which utilizes the variance that is explainable in principle, Var_exp_[*r*] (Eq. 6), and the variance of the residuals corrected by the observation noise,

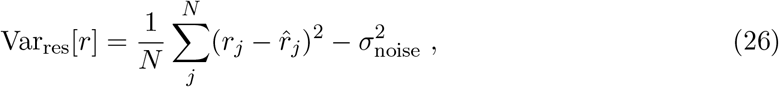

where *j* indexes images. This measure corrects for observation noise, which variance 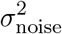 we estimated with Eq. (7). To compute model performance we averaged the FEV across neurons.

### 4.7 In silico experiments

To perform cross-orientation and size tuning experiments *in silico*, we first had to determine the optimal Gabor pattern that elicited maximum response for all units in the models we investigated. We specified an image containing a Gabor pattern by the Gabor’s center location, size, spatial frequency, phase, orientation and contrast. For each neuron and model, we obtained the optimal Gabor image by finding the parameter combination that lead to maximum model prediction. For the center location we tested all positions in the 40 px× 40 px (1.14° of visual field) image using 12 different orientations with equally spaced angles *ϕ/π* = 0, 1*/*12, 2*/*12, …, 11*/*12 (in units of *π*) and 8 phasesψ*/*2*π* = 0, 1*/*8, 2*/*8, …, 7*/*8 (in units of 2*π*). We searched for size, spatial frequency and contrast with equidistant values in log-space. Specifically, we sampled 8 sizes *s*_*i*_ = 4 px · (1.3895)^*i*^, *i* = 0, 1, …, 7, resulting in a maximum size of 40 px, where the size of the Gabor is defined as ±2 standard deviations of its Gaussian envelope. For spatial frequency, we used 10 values *f*_*i*_ = (1.3)^−1^ · (1.3)^*i*^, *i* = 0, 1, 2, …, 9 measured in cycles per four standard deviations of the according Gaussian envelope. We determined the maximum contrast from the images in our training set. After normalization, the mean across all pixel values was zero, the minimum and maximum pixel values were −2.52 and 3.20, respectively. Thus, making sure that the Gabor’s pixel values are within the value range seen by the models during training, we set the maximum contrast to *c*_max_ = 2 · 2.52 corresponding to a maximum amplitude of *a*_max_ = 2.52. We chose 6 amplitude (contrast) values *a*_*i*_ = *c*_*i*_*/*2 = 0.01 *a*_max_ · (2.51189)^*i*^, *i* = 0, 1, 2, …, 5 resulting in *c*_5_ = *c*_max_.

#### 4.7.1 Cross-orientation inhibition

For the cross-orientation inhibition experiment, we used the parameters determined for the optimal Gabor to generate plaids (Fig. 4A, insets) by linear superposition of the optimal Gabor with a 90° rotated version of it. We sampled the contrasts of both Gabor components from all combinations out of 𝒞 = {0, *c*_*i*_} with *c*_*i*_ = 0.01 *c*_max_ · (1.63069)^*i*^, *i* = 0, 1, 2, …, 8 where the highest contrast was 50% of the maximum contrast, *c*_8_ = 0.5 *c*_max_, making sure the pixel values of the combined plaid stimulus stayed within the value range seen by our models during training. In addition, for the orthogonal Gabor we used 8 phasesψ*/*2*π* = 0, 1*/*8, 2*/*8, …, 7*/*8 (in units of 2*π*). We obtained contrast tuning curves presenting all plaid stimuli averaging across phases to get results that are independent of relative phase between optimally and orthogonally oriented Gabor components. To quantify cross-orientation inhibition strength, we defined a cross-orientation inhibition index (COI) measuring the relative reduction in response due to adding an orthogonal Gabor with contrast *c*_*i*_ to the driving Gabor of optimal orientation with contrast *c*_*j*_ for all contrast combinations (*c*_*i*_, *c*_*j*_) ∈ 𝒞_orthogonal_ × 𝒞_optimal_,

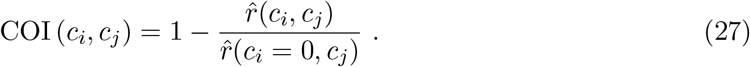

We define a cell to be cross-orientation inhibited if the cross-orientation inhibition index COI is at least 10% for any combination of optimal and orthogonal Gabor contrasts, i.e. the presence of the orthogonal Gabor reduces response by at least 10%.

#### 4.7.2 Size tuning and surround-suppression control analysis

To obtain size tuning curves, we generated gratings using the parameters of the optimal Gabor for each cell. Then, instead of presenting a Gabor with a Gaussian envelope, we present a sinusoidal grating with full contrast and a sharp cut-off to zero intensity (50% grey) outside of a circular area with predefined diameter (Fig. 7). We set the maximum grating diameter to *d*_max_ = 120 px (3.43° of visual field), making sure to cover the whole 40 px × 40 px (1.14° of visual field) image even if the grating’s center would be in one of the image corners. Then, we chose diameters *d*_*i*_ = 0.05 *d*_max_ · (1.23859)^*i*^, *i* = 0, 1, 2, …, 14. As almost all optimal Gabors had maximum contrast, we presented the gratings in this experiment with maximum contrast, too. To quantify any surround suppression, we used the suppression index (SI) proposed by Cavanaugh et al. (2002a), measuring the reduction from the highest model prediction 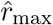 to the prediction 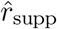 for a grating completely filling the whole image for each neuron,

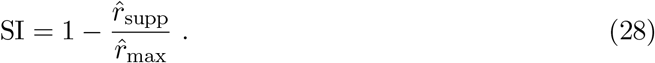

Hence, a SI of zero corresponds to no suppression, whereas a fully suppressed cell would lead to a SI of one.

### 4.8 Evaluation of orientation-specific normalization

To analyse how the preferred orientation of the features being normalized depend on that of the features providing normalizing inputs (Fig. 5), we determined for each feature map whether it extracts oriented features and – if so – its preferred orientation. To do so, we windowed each convolutional kernel of size 13 px × 13 px (0.37° of visual field) with a Gaussian window (standard deviation: 3 px, corresponding to 0.09°), normalized it and then computed its 2D power spectrum (using the discrete Fourier transform with 64 × 64 samples). We then quantified how power spectral density is distributed across orientations by computing a mean resultant vector *m* given by:

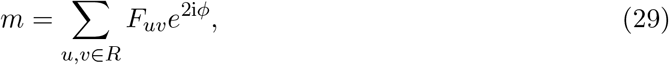

where *F*_*uv*_ is the Fourier transformed kernel, 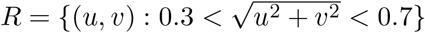 contains all frequencies between 0.3 and 0.7 (with 1.0 being the Nyquist frequency), *ϕ* = atan2(*v, u*) is the orientation, i the imaginary unit and the factor 2 in the complex exponential accounts for the fact that we are interested in orientation, which is periodic in 180° or *π*. If all power in a kernel is concentrated in one orientation, the mean resultant vector will be long, whereas an unoriented kernel will have a mean resultant vector near zero. Based on visual inspection of the kernels in model fits of the top 11 to 20 model runs (in terms of validation set accuracy), we found |*m*| = 0.125 to be a reasonable threshold for separating oriented from unoriented features and used it as a heuristic for further analyses of the best ten model runs, avoiding issues with post-hoc statistical testing.

To quantify how strong a feature *l* is normalized by other features *k*, we computed the average normalizing input, which is given as the expected value (over images) of the product *p*_*kl*_ · *y*_*kuv*_(*x*) in Eq. (1). Since this normalization input depends on the stimulus, we computed its expected value of all images in the validation set. We removed the dependence on space by averaging over all locations within the feature map.

### 4.9 Control: All channels contribute to our model’s prediction

To verify that all features contribute to normalization, we analyzed the readout feature weights for the best ten models (assessed in terms of performance on the validation set). However, there are two issues that prevent a direct comparison across models and neurons of the feature weightings. First, the factorization of the readout into spatial and feature weightings is not unique: scaling the spatial weights (*a* in Eq. (11)) by a factor *β* whilst scaling the feature weights (*b* in Eq. (11)) by 1*/β* yields the same output limiting comparisons across neurons. Second, a similar exercise between the normalization weights *p* and the semi-saturation constant *σ* (Eq. 10) impedes comparison across models. To solve these issues, we normalized the feature readout weights of each neuron for this control analysis so that the resulting vectors for each neuron and model convey how much a certain feature map contributes to predict a neuron’s response compared to the other feature maps, making the feature readout weights comparable across neurons. Next, we averaged these weights across neurons to assess the importance of the features in a model. Since these normalized feature readout weights were comparable across both neurons and models, we calculated a collective distribution of the averaged feature readout weights from the best ten models. To make sense of this distribution’s absolute values, we evaluated its width in terms of the coefficient of variation, which is the standard deviation in units of the mean.

### 4.10 Implementation details

We used Tensorflow (Abadi et al., 2015) to implement models as well as Python, which we additionally used for data analysis. We optimized models with the Adam optimizer (Kingma and Ba, 2015) using mini-batches of size 256. In addition, we used the Python packages Numpy/Scipy (Walt et al., 2011), Pandas (McKinney, 2010), Matplotlib (Hunter, 2007), Seaborn (Waskom et al., 2017), Datajoint (Yatsenko et al., 2015, 2018) and the tools Jupyter (Kluyver et al., 2016) and Docker (Merkel, 2014).

## Acknowledgements

We thank Fabian H. Sinz, Felix A. Wichmann, David A. Klindt, Alexander Böttcher, Matthias Kümmerer, Dylan M. Paiton, Robert Geirhos, Claudio Michaelis and Ivan Ustyuzhaninov for valuable discussions. M.F.B. and S.A.C. thank the International Max Planck Research School for Intelligent Systems (IMPRS-IS). The research was supported by the German Federal Ministry of Education and Research (BMBF) via the Competence Center for Machine Learning (FKZ 01IS18039A); the German Research Foundation (DFG) grant EC 479/1-1 (A.S.E.), the Collaborative Research Center (SFB 1233, Robust Vision) and the Cluster of Excellence “Machine Learning – New Perspectives for Science” (EXC 2064/1, project number 390727645); the Bernstein Center for Computational Neuroscience (FKZ 01GQ1002); the National Eye Institute of the National Institutes of Health under Award Numbers R01EY026927 (A.S.T.), DP1 EY023176 (A.S.T.), and NIH-Pioneer award DP1-OD008301 (A.S.T). This research was also supported by NEI/NIH Core Grant for Vision Research (EY-002520-37), NEI training grant T32EY00700140 (G.H.D) and F30EY025510 (E.Y.W.), DARPA grant N66001-17-C-4002, and the Intelligence Advanced Research Projects Activity (IARPA) via Department of Interior/Interior Business Center (DoI/IBC) contract number D16PC00003. The U.S. Government is authorized to reproduce and distribute reprints for Governmental purposes notwithstanding any copyright annotation thereon. Disclaimer: The views and conclusions contained herein are those of the authors and should not be interpreted as necessarily representing the official policies or endorsements, either expressed or implied, of IARPA, DoI/IBC, or the U.S. Government. The funders had no role in study design, data collection and analysis, decision to publish, or preparation of the manuscript.

## Data availability

The data analyzed in this study is available online at https://doi.org/10.12751/g-node. 2e31e3. The code that supports the findings of this study will be made available online upon publication.

